# Why detailed modelling matters in the pre-clinical evaluation of temporomandibular joint implants

**DOI:** 10.1101/2025.04.28.651047

**Authors:** Girish Chandra, Rajdeep Ghosh, Vivek Verma, Kamalpreet Kaur, Ajoy Roychoudhury, Sudipto Mukherjee, Anoop Chawla, Kaushik Mukherjee

**Author notes:** Girish Chandra and Rajdeep Ghosh are designated as joint first authors. Corresponding author: Dr. Kaushik Mukherjee Department of Mechanical Engineering, Indian Institute of Technology Delhi New Delhi, 110 016.

## Abstract

End-stage temporomandibular joint (TMJ) disorders often necessitate joint replacement to restore bilateral mastication. Patient-specific TMJ implants typically rely on detailed in silico modelling derived from subject-specific computed tomography (CT) scans of the mandible. However, generating highly detailed finite element (FE) models is computationally expensive. Although structural simplifications of the mandible are known to influence mechanical behaviour, their quantitative impact on TMJ implant evaluation has not been systematically investigated. This study examines the influence of three modelling simplifications on stress–strain predictions in intact and implanted mandibles: (i) a detailed segmented mandible with tissue-specific material properties (Model 1), (ii) a simplified mandible comprising cortical bone only (Model 2), and (iii) a further simplified mandible excluding dental crowns (Model 3). Both narrow and standard TMJ implants were analysed under osseointegrated and non-osseointegrated conditions, resulting in fifteen FE models. Physiological clenching activities were simulated. Compared with the detailed model, simplified models showed reductions of up to 50% in maximum principal strain in bone and up to 44% in von Mises stress in TMJ implants, while preserving spatial stress–strain trends. These findings indicate that simplified models may be suitable for preliminary implant design, whereas detailed FE modelling remains essential for final pre-clinical evaluation.

## Introduction

The temporomandibular joint facilitates essential functions, including mouth opening/closing, clenching/biting, and speaking. In a healthy human, these masticatory activities can go up to 1 million times annually (Nakhaei et al., 2023). In addition, the mandible has several important functions, being directly related to respiration, enunciation, and deglutition, and is also related to an individual’s aesthetic (Al-Ahmari et al., 2015; Kim and Donoff, 1992; Luo et al., 2017; Seol et al., 2014). However, approximately 5% of the human population experiences limitations in these activities due to end-stage TMJ disorders such as ankylosis (Balon et al., 2019; Ingawalé and Goswami, 2009). Total joint replacement (TJR) using TMJ implants is typically done to treat these conditions (Gonzalez-Perez et al., 2020; Yoda et al., 2020).

With the recent advancement in 3D printing and rapid prototyping (Ackland et al., 2017), the latest focus is on the design and development of patient-specific TMJ implants (Ingawale and Goswami, 2022). Designing patient-specific TMJ implants warrants detailed *in silico* evaluation to estimate post-operative biomechanical performances of these implants under masticatory loading (Banerjee et al., 2023; Deshmukh et al., 2012; Ghosh et al., 2024; Ingawale and Goswami, 2022; Kashi et al., 2010; Pinheiro et al., 2021; Vignesh et al., 2020). While some of these studies (Ghosh et al., 2024; Pinheiro et al., 2021) developed detailed FE models of the mandible, with components like enamel, dentin, periodontal ligament (PDL), cancellous and cortical region, and analyzed the implant performance under masticatory loading, there are numerous studies which simplified the FE model of mandible either by neglecting dentures (Huang et al., 2015; Oguz et al., 2009) or by considering the entire mandible to be made of cortical bone (Filardi, 2020; Huang et al., 2021; Kharmanda and Kharma, 2017; Mesnard et al., 2011; Wang et al., 2017). Although these types of mandible simplifications save time and effort associated with segmenting a clinical CT dataset, such simplifications do alter the FE predicted results. However, there is no study reporting the extent of influence of such modelling simplifications on the FE-based design evaluation of TMJ implants.

Hence, this study investigates the extent to which such geometric and material simplifications could influence the pre-clinical evaluation of mandible and TMJ implants by considering the following three model simplifications for intact and implanted mandibles: a detailed segmented mandible with tissue-specific material properties (Stage 0 simplification, Model 1), a simplified mandible, derived from Model 1, made of cortical bone only (Stage I simplification, Model 2), and a further simplified mandible, derived from Model 2, wherein the dental crowns are excluded (Stage II simplification, Model 3). For implanted mandible models, narrow and standard TMJ implants (45 mm long, Zimmer Biomet Microfixation system, Jacksonville, FL, USA) were considered under both osseointegrated and non-osseointegrated conditions. Thus, a total of 15 FE models were developed (out of which three were intact mandibles, and 12 were implanted mandibles considering all combinations of three levels of simplifications, two types of TMJ implants, two interfacial conditions i.e. osseointegrated and non-osseointegrated) to understand the influence of modelling simplifications on stresses and strains in mandible and TMJ implants (mandibular component and screws) under masticatory loading.

## Materials and Methods

Clinical CT images (512 x 512 pixels; pixel size of 0.439 mm; slice thickness of 0.75 mm) of a healthy anonymized 20-years-old male subject were acquired. Necessary ethics approval for this study was provided by the Institute Ethics Committee of Centre for Dental Education and Research, AIIMS, New Delhi, India (IEC-611/15.07.2022, AA-3/05.08.2022, RP-11/2022). CT-specific mandible was developed in Materialise Mimics v25.0 (Materialise Inc., Leuven, Belgium) and meshed in Hypermesh v2022 (Altair Engineering Inc., Troy, MI, USA). Further FE-based biomechanical evaluations were performed in ANSYS Mechanical APDL v2022 R2 (ANSYS Inc., Canonsburg, PA, USA). Details of the FE model development process are being presented in subsequent sections.

### Development of Intact Mandible Model

The subject-specific, detailed, intact mandible model (Stage 0 simplification, Model 1, Figure 1-b&c) was from our previous study (Ghosh et al., 2024). The model had multiple components, including cortical bone, cancellous bone, teeth, PDL, articular fibrocartilage, and temporal bone blocks. Each component was assigned tissue-specific material properties (Ghosh et al., 2024). Further details of the model generation procedure, including detailed verification and validation of the FE model, are described in our previous study (Ghosh et al., 2024).

**Figure 1.**
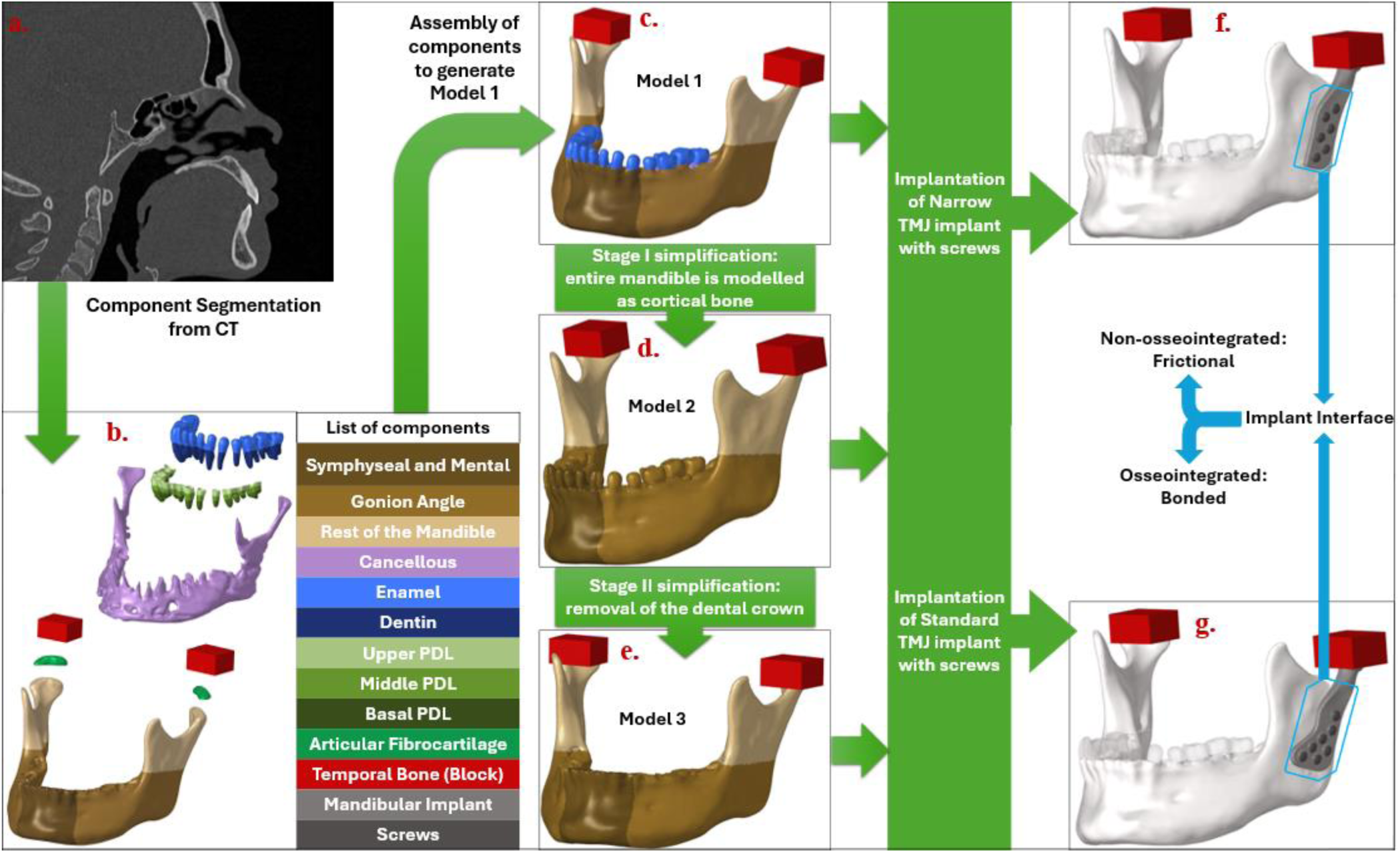
Overview of the methodology: (a) CT data from an anonymous subject, (b) exploded view of detailed intact mandible model (Model 1), (c) detailed intact mandible model (Model 1) (d) simplified intact mandible model (Model 2) developed from Model 1, (e) simplified intact mandible model (Model 3) without crown of teeth (f) osteotomised mandible with transparent teeth including narrow TMJ implants showing implant interface region, and (g) osteotomised mandible with transparent teeth including standard TMJ implants showing implant interface region.

A simplified model of an intact mandible (Stage I simplification, Model 2, Figure 1-d) was derived from this detailed mandible by modelling the entire mandible (cortical bone, cancellous bone, PDL and teeth) as cortical bone. The model, however, had articular fibrocartilage on its condyle and the temporal bone blocks. A further simplified model of the intact mandible (Stage II simplification, Model 3, Figure 1-e) was derived from Model 2 by excluding the dental crowns. All these mandibular models were discretised with quadratic tetrahedral elements with edge lengths of 0.3 – 2 mm based on the mesh convergence study performed earlier (Ghosh et al., 2024).

### Development of Implanted Mandible Models

The 3D models of Biomet narrow and standard mandibular stock implant components of size 45 mm (Zimmer Biomet Microfixation system, Jacksonville, FL, USA; Figure 1 - f & g) were obtained from micro-CT imaging (Xtreme CT II, Scanco Medical, Switzerland, isotropic voxel size: 60 μm). Based on Biomet mandibular screws (BIOMET Microfixation, 2007), bi-cortical screws were modelled using SolidWorks 2022 (Dassault Systems, Vélizy-Villacoublay, France) as cylinders (Φ2.7 mm, length: 8 mm – 12 mm). The models of the intact mandible and TMJ implant were assembled into Rhinoceros^®^ (Rhinoceros V7.0, Robert McNeel & Associates, Seattle, USA) for virtual implantation based on clinical recommendations. Based on our previous study (Ghosh et al., 2024), quadratic tetrahedral elements with mesh size of ∼ 0.5 mm edge length were used to discretise the TMJ implants. For each intact mandible model, corresponding implanted models were developed with two types of stock TMJ implants (Narrow and Standard).

### Interfacial conditions

This study considered both the immediate post-operative non-osseointegrated and the long-term osseointegrated conditions. To simulate non-osseointegrated (NOI) condition, a frictional (µ= 0.3) implant-bone interface was modelled, whereas for osseointegrated (OI) interface condition, a bonded bone-implant interface was considered between the mandibular component and the cortical bone. Thus, finally, a total of 12 implanted mandible models were being generated, out of which four models were based on Model 1 and were from our previous study (Ghosh et al., 2025), the next four models were based on Model 2 (stage I simplification) and the next four models were based on Model 3 (stage II simplification).

A frictionless interface was modelled between the fibrocartilage layer and the mandibular condyle in the intact mandibles. The interface between the head (uppermost region) of the mandibular component (TMJ implant) and the rigid temporal bone block (as fossa component) was, however, considered frictional (µ = 0.3) in the implanted mandible (Ghosh et al., 2024). All other interfaces (implant-screw and mandible-screw) were considered bonded. The developed FE model was solved using augmented Lagrangian algorithm.

### Material properties

In Model 1, tissue-specific linear elastic material properties (Table 1) were assigned to each anatomical segment: cortical (region-specific orthotropic), cancellous, PDL (region-specific isotropic), and teeth (region-specific isotropic)(Ghosh et al., 2025; Korioth et al., 1992). In the simplified models (Model 2 and Model 3), region-specific cortical bone material property (Korioth et al., 1992) was prescribed to the mandible bone. For all models, the articular fibrocartilage, and implants were modelled using the isotropic linear elastic material properties, while the temporal bone block was considered rigid (Table 1). TMJ implants (mandibular component and screws) were assumed to be made of Ti-alloy (Ti-6Al-4V) Medical Grade 23 ELI (Table 1).

**Table 1.** Material properties of the components of the FE models (Korioth et al., 1992)

| Components | Young's Modulus (GPa) |  |  | Poisson's Ratio |  |  | Shear Modulus (GPa) |  |  |
| --- | --- | --- | --- | --- | --- | --- | --- | --- | --- |
| | $E_x$ | $E_y$ | $E_z$ | $\nu_{xy}$ | $\nu_{yz}$ | $\nu_{xz}$ | $G_{xy}$ | $G_{yz}$ | $G_{xz}$ |
| Cortical |  |  |  |  |  |  |  |  |  |
| Symphyseal and mental | 23 | 10 | 15 | 0.3 | 0.3 | 0.3 | 4.8 | 3.6 | 6.2 |
| Gonial angle | 20 | 11 | 12 | 0.3 | 0.3 | 0.3 | 4.8 | 5.3 | 6.0 |
| Rest | 17 | 6.9 | 8.2 | 0.3 | 0.3 | 0.3 | 2.8 | 2.9 | 4.6 |
| Cancellous | 0.96 | 0.32 | 0.39 | 0.3 | 0.3 | 0.3 | 0.09 | 0.13 | 0.17 |
| Enamel | 80 | - | - | 0.3 | - | - | - | - | - |
| Dentin | 17.6 | - | - | 0.25 | - | - | - | - | - |
| Periodontal ligament |  |  |  |  |  |  |  |  |  |
| Coronal third | 0.0032 | - | - | 0.45 | - | - | - | - | - |
| Middle third | 0.0027 | - | - | 0.45 | - | - | - | - | - |
| Apical third | 0.0025 | - | - | 0.45 | - | - | - | - | - |
| Articular Fibrocartilage | 0.006 |  |  | 0.47 |  |  |  |  | - |
| Mandibular component/screws (Ti-6Al-4V: ELI) | 113.8 |  |  | 0.342 |  |  |  |  | - |
| Block (Temporal Bone) | Rigid |  |  | Rigid |  |  |  |  | - |

### Loading and Boundary Conditions

Six clenching load cases (Supplementary Table 1) were considered in the present study to simulate a complete mastication cycle: incisal (INC), intercuspal (ICP), right molar (RMOL), left molar (LMOL), right group (RGF), and left group (LGF) bites (Ghosh et al., 2024), (Dutta et al., 2023; Hijazi et al., 2016). These tasks primarily fall into three clenching categories: central (INC), unilateral (RMOL, LMOL, RGF, and LGF), and bilateral (ICP). Muscle forces from the following seven muscles (Supplementary Figure 1)(Hijazi et al., 2016) were considered in the present study: two masseters (superior masseter and deep masseter), two pterygoids (medial pterygoid/lateral pterygoid), and three temporalis (anterior temporalis/middle temporalis/posterior temporalis).

However, the effect of the lateral pterygoid on the implanted side was ignored due to the removal of the condyle (attachment site of lateral pterygoid). Furthermore, the current study also hypothesized the effective reattachment of all other muscles post-surgery. The top surface of *os temporale* was fully constrained (Hijazi et al., 2016; Korioth et al., 1992). The displacements at bite points (the uppermost region on teeth) corresponding to each clenching task were restrained in Z-direction (Pinheiro et al., 2021).

### Interpretation of FE results

Previous studies considered peak principal strains as a failure criterion for brittle cortical bones. (Fan et al., 2023; Schileo et al., 2008). Consequently, in the current investigation, peak principal strains in the bone were used to compare the biomechanical behaviour of intact and reconstructed mandibles. For comparing the behaviour of ductile TMJ implants, von Mises stress, however, was used as a failure criterion. To avoid strain and stress singularity in the FE models, 99^th^ percentile values were considered for comparison(Ghosh et al., 2025). FE results from the simplified models (Models 2 and 3) were compared with those of the detailed mandible model (Model 1) using linear regression and the coefficient of determination (R²) for ICP case. Principal strains in the mandibular cortex and von Mises stress in the implant and screws were considered for this comparison.

## Results

A thorough comparison of stresses and strains in implants and bone under different clenching activities is presented here. First, the principal strain distributions across the bone were analysed for intact and reconstructed mandibles, and, thereafter, the von Mises stresses in mandibular components and screws were analysed to investigate the influence of different modelling simplifications on FE-based preclinical evaluations. Since the ICP loading condition induced maximum stresses and strains in the mandible, implant and screws (Ghosh et al., 2024), corresponding distributions are included in the main manuscript, while stress-strain distributions for other loading conditions are included in the supplementary file (Supplementary Figures 2 and 3).

### Biomechanical evaluation of the mandible

Maximum principal strains in the intact and implanted mandible ranged between 394 – 1880 με (Figure 2) for different loading conditions under different modelling simplifications. Among all the clenching tasks, the ICP load case resulted in the highest maximum principal strain (1880 με), the RGF/LGF load cases induced the lowest maximum principal strain (394 με) for all three modelling simplifications. For all the loading conditions, Model 1, with the detailed mandible, exhibited the highest maximum principal strains in bone (range: 610 - 1880 με). In comparison, Model 2 exhibited the least maximum principal strains (range: 394 - 994 με), whereas strains in Model 3 were intermediate (range: 420 - 1040 με). Upon comparing the maximum principal strains in the mandible between non-osseointegrated (immediate post-operative) and osseointegrated (long-term) conditions, it was observed that the osseointegration-induced strain reduction in the mandible was captured by all the modelling simplifications. However, the extent of strain reduction was highly influenced by the modelling simplifications.

**Figure 2.**
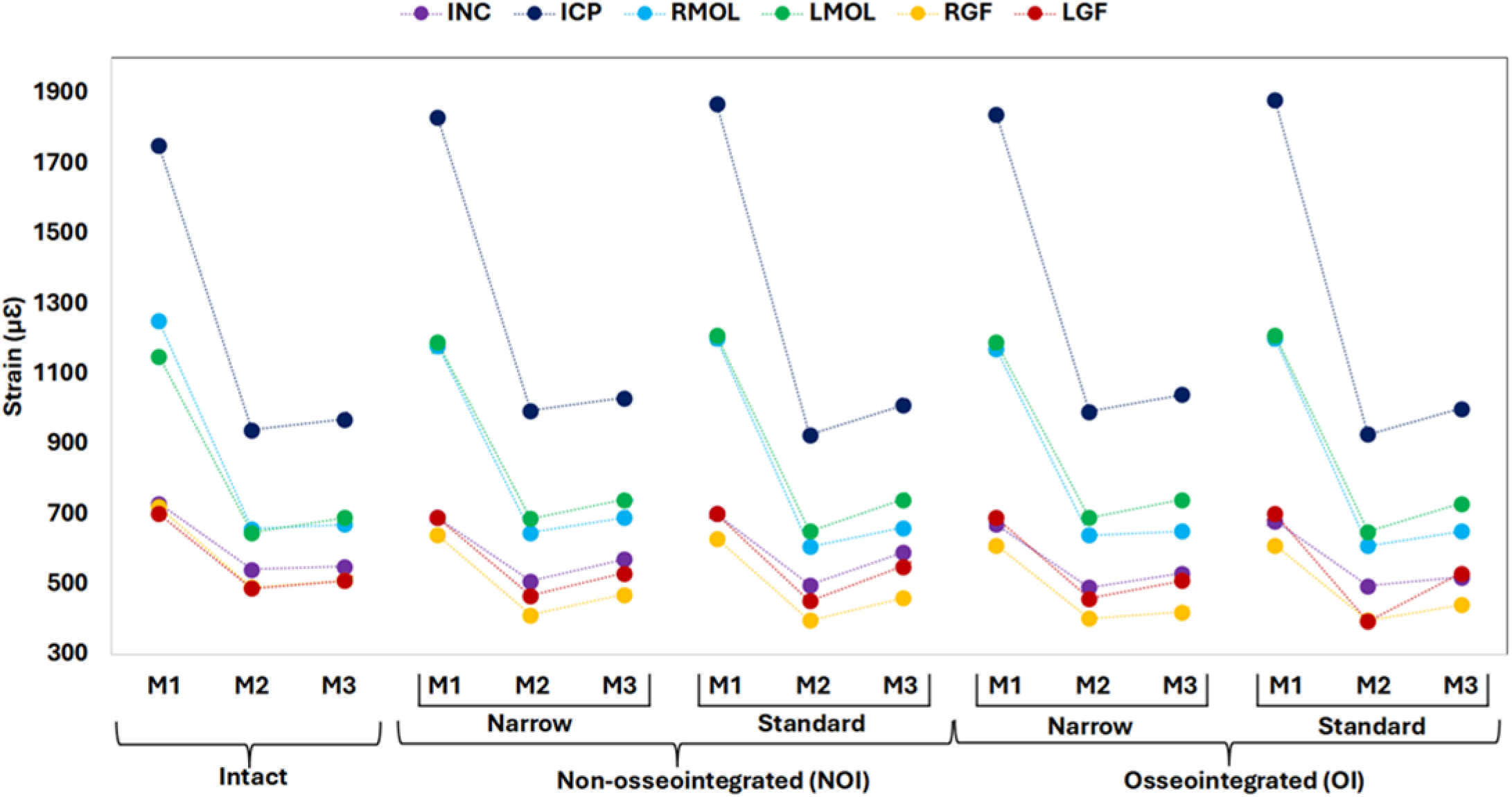
Variation in maximum principal strains in the intact and implanted (with narrow/standard mandibular component) mandible across six clenching tasks (under complete mastication cycle) in Model 1 (M1), Model 2 (M2) and Model 3 (M3).

The maximum principal strain distribution in intact as well as implanted mandibles (with narrow and standard mandibular components) under the ICP clenching task (Figure 3) exhibited how the predicted strains in the mandibles reduced with simplifications. For intact mandible, regions of low strains were observed in the symphyseal and mental regions, irrespective of the loading and modelling techniques used. However, Models 2 and 3 exhibited higher areas of low strains (marked in yellow) as compared to Model 1. For the implanted mandible models with standard TMJ implants, lower strains around the coronoid process were observed in Models 2 and 3 (marked in black) as compared to Model 1(marked in red). No such visible difference in maximum principal strain distribution was, however, there for the narrow implant for different modelling simplifications. High strains near the screw holes were observed in non-osseointegrated condition, as compared to the osseointegrated condition, for all the modelling simplifications.

**Figure 3.**
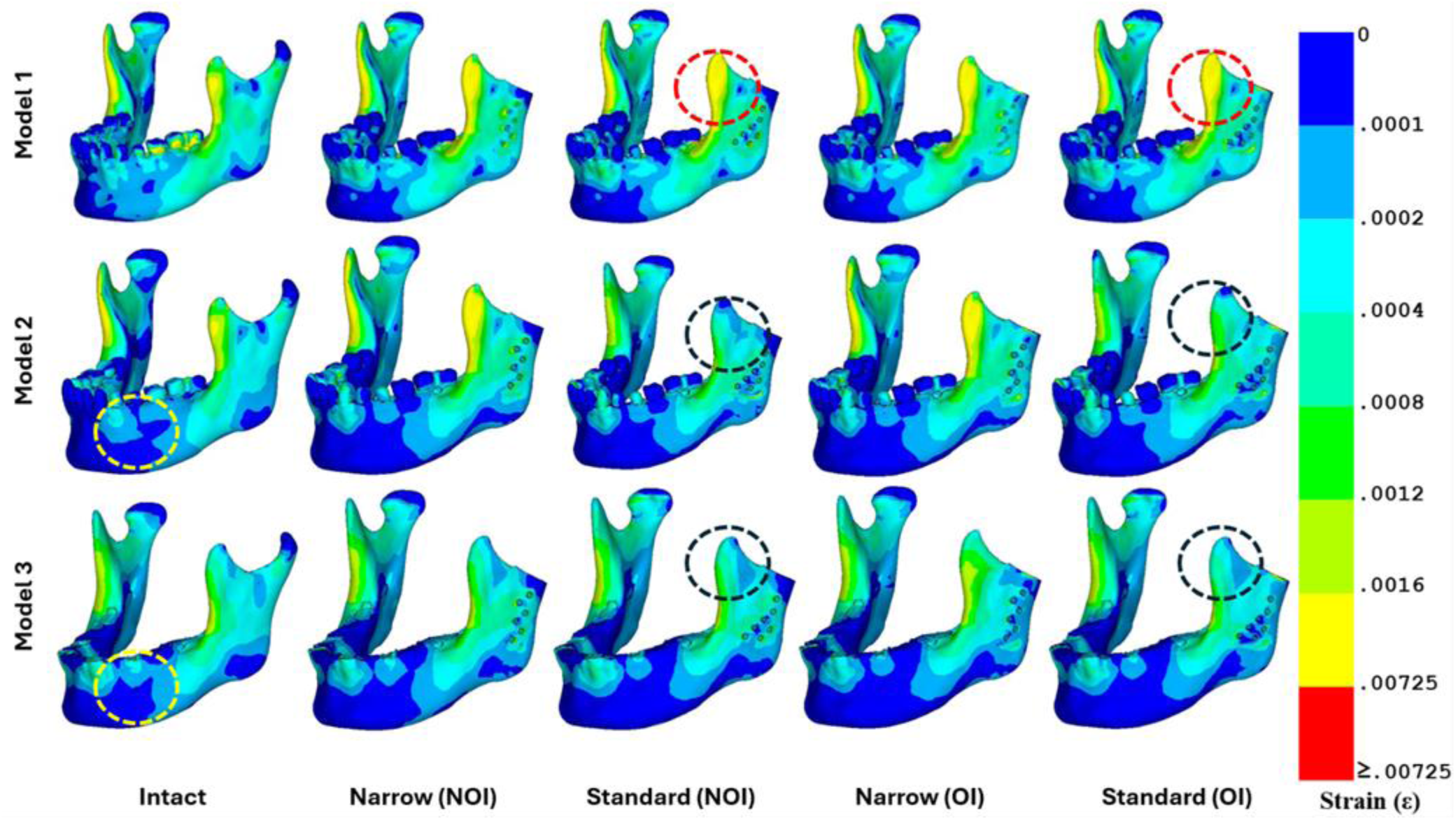
Comparison of maximum principal strain distribution on the cortical bone in various FE models (Model 1: Detailed Segmented Mandible, Model 2: Cortical Mandible, Model 3: Cortical mandible without crown of teeth, NOI: non-osseointegrated, OI: osseointegrated) under ICP bite.

### Stresses in TMJ Implants

Figure 4 shows the magnitude of von Mises stress in mandibular components and screws of TMJ implants. Overall, ICP load induced the highest stresses in narrow and standard TMJ implants and screws, whereas LMOL caused the lowest stresses (Figure 4). However, for all clenching tasks, the von Mises stresses in TMJ implants and screws were much less than the yield strength (∼800 MPa) of Ti-alloy considered in this study (Farooq et al., 2020). The three modelling simplifications were found to alter the von Mises stresses in implants and screws, depending upon the level of simplification made to develop those biomechanical models. Overall, the maximum von Mises stresses in the TMJ implant in Model 1 were higher than those in Model 2 which, in turn, was higher than those in Model 3 (Figure 4). Similar trends were also observed for the screws (Figure 4). A comparison between osseointegrated and non-osseointegrated conditions revealed that for all three modelling simplifications, osseointegration causes a reduction in stresses in implants and screws (Figure 4)

**Figure 4.**
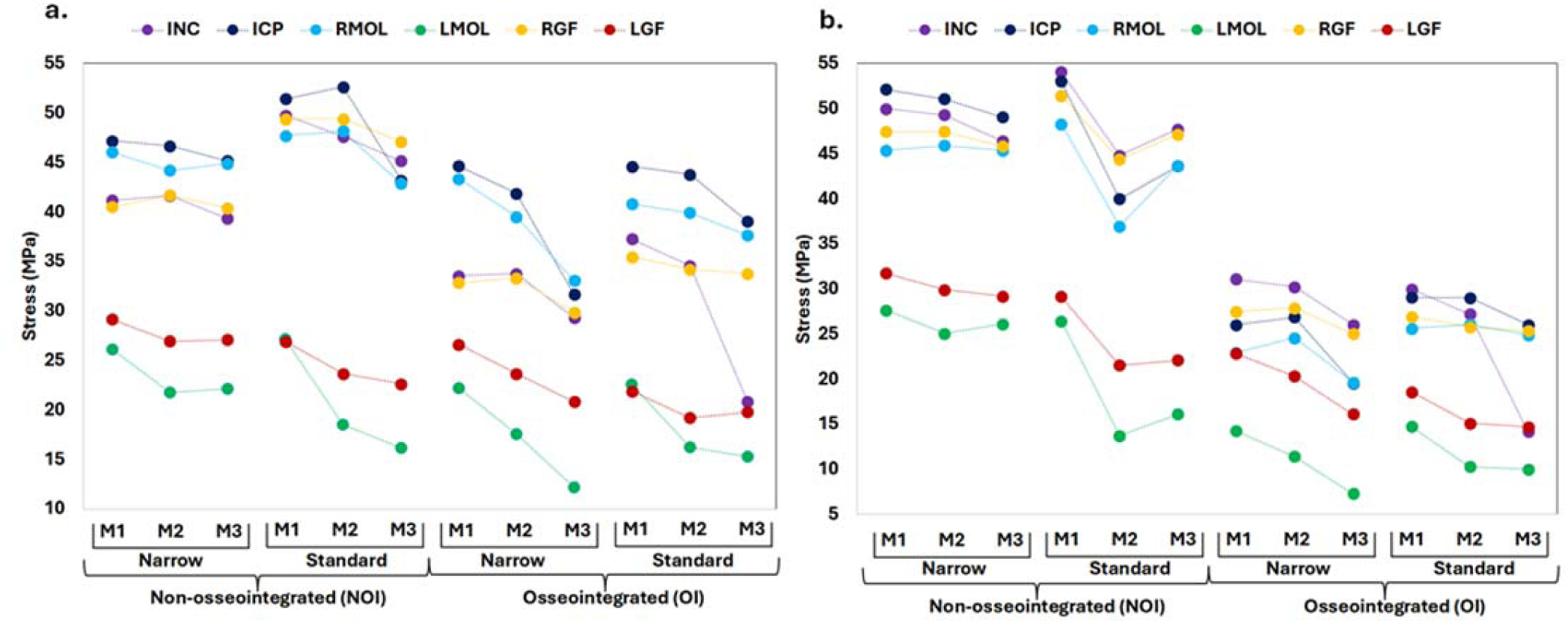
Variation in von Mises stresses in (a) mandibular components and (b) screws across six clenching tasks in Model 1 17, Model 2 and Model 3.

Figure 5 shows the equivalent stress distribution in mandibular components and screws of TMJ implants for ICP load case. A marked difference in stress distribution was observed between the narrow and the standard implant. Lower stress was observed at the inferior lateral flares of the standard implant (as marked in magenta in Figure 5), unlike the narrow ones, which show an elevated stress at the inferior implant regions (marked in red).

**Figure 5.**
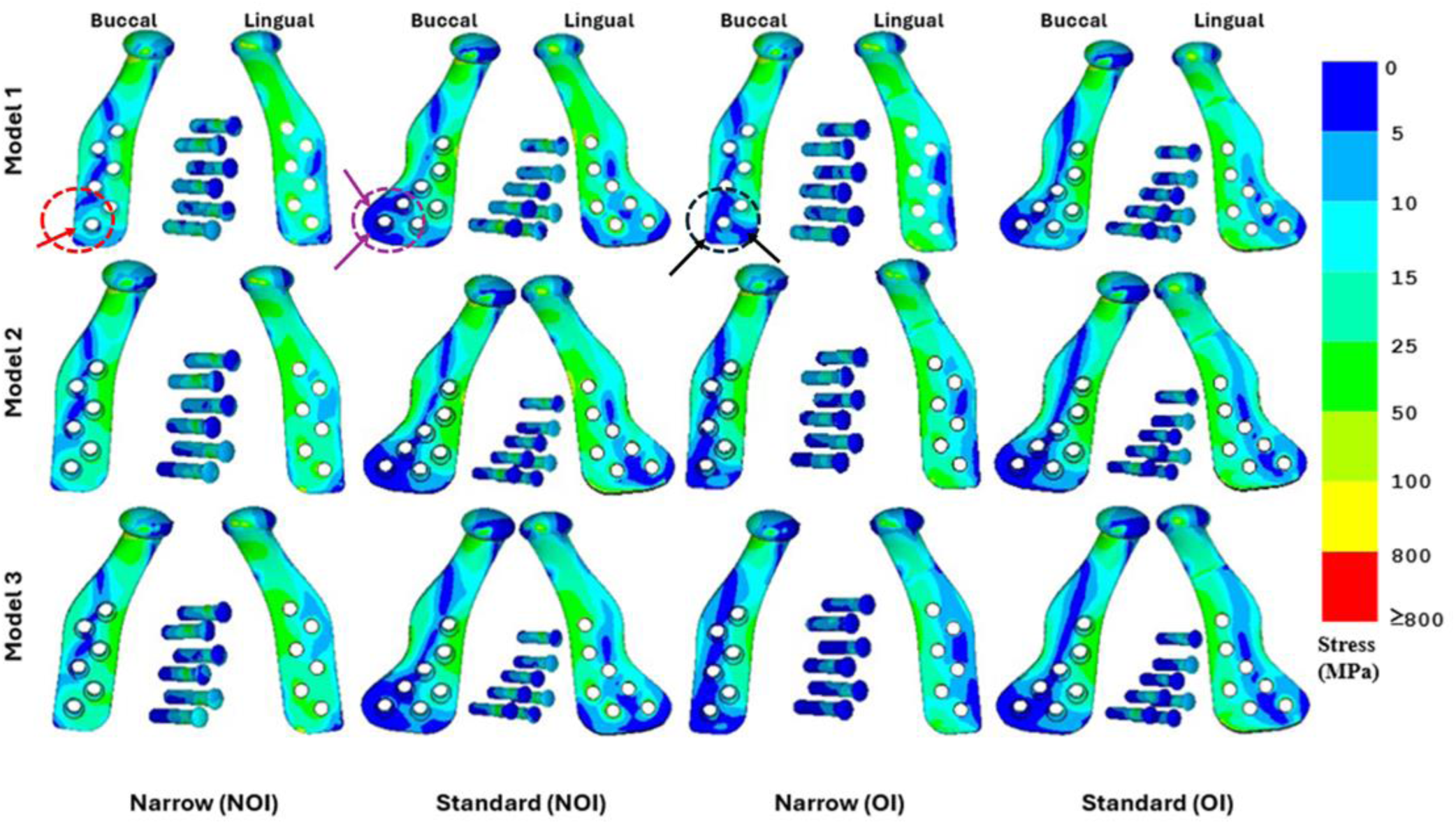
Comparison of von Mises stress distribution on the implants (Mandibular component and Screws) in various FE models (Model 1: Detailed Segmented Mandible, Model 2: Cortical Mandible, Model 3: Cortical mandible without crown of teeth, NOI: non-osseointegrated, OI: osseointegrated) under ICP bite. The legends were chosen based on the failure limit of Ti6Al4V medical grade 23(yield strength=800MPa) ^33^.

Moreover, the buccal side, which was geometrically far from the bone, experienced more low-stress regions compared to the lingual side in both implants. However, when comparing the state of osseointegrations, osseointegrated states experience much lower stress in both implants, with a visibly marked difference in stress in the inferior regions of narrow implants (marked in black). Lower stress regions were also observed in screws in their osseointegrated state as compared to their initial non-osseointegrated state. Higher stresses were consistently observed near the screw holes in both the implants and the screws. Similar spatial stress distribution patterns across the implants and interface conditions were observed when comparing all three modelling techniques.

In addition to comparing the von Mises stresses of the mandibular components and screws between both interfaces and both types of implants, it was found that all modelling simplifications resulted in overall stress reductions in TMJ implants (ranging from 2.4% to 44.1%) and screws (ranging from 0.1% to 48.8%) across all loading conditions.

### Statistical relationship between the mechanical metrics obtained with different modelling techniques

For further quantitative comparison, the location-specific stresses and strains in corresponding elements of the mandibular cortex, TMJ implant, and screws were compared for ICP case between the detailed model (Model 1) and simplified models (Models 2 and 3), as shown in Figure 6.

**Figure 6.**
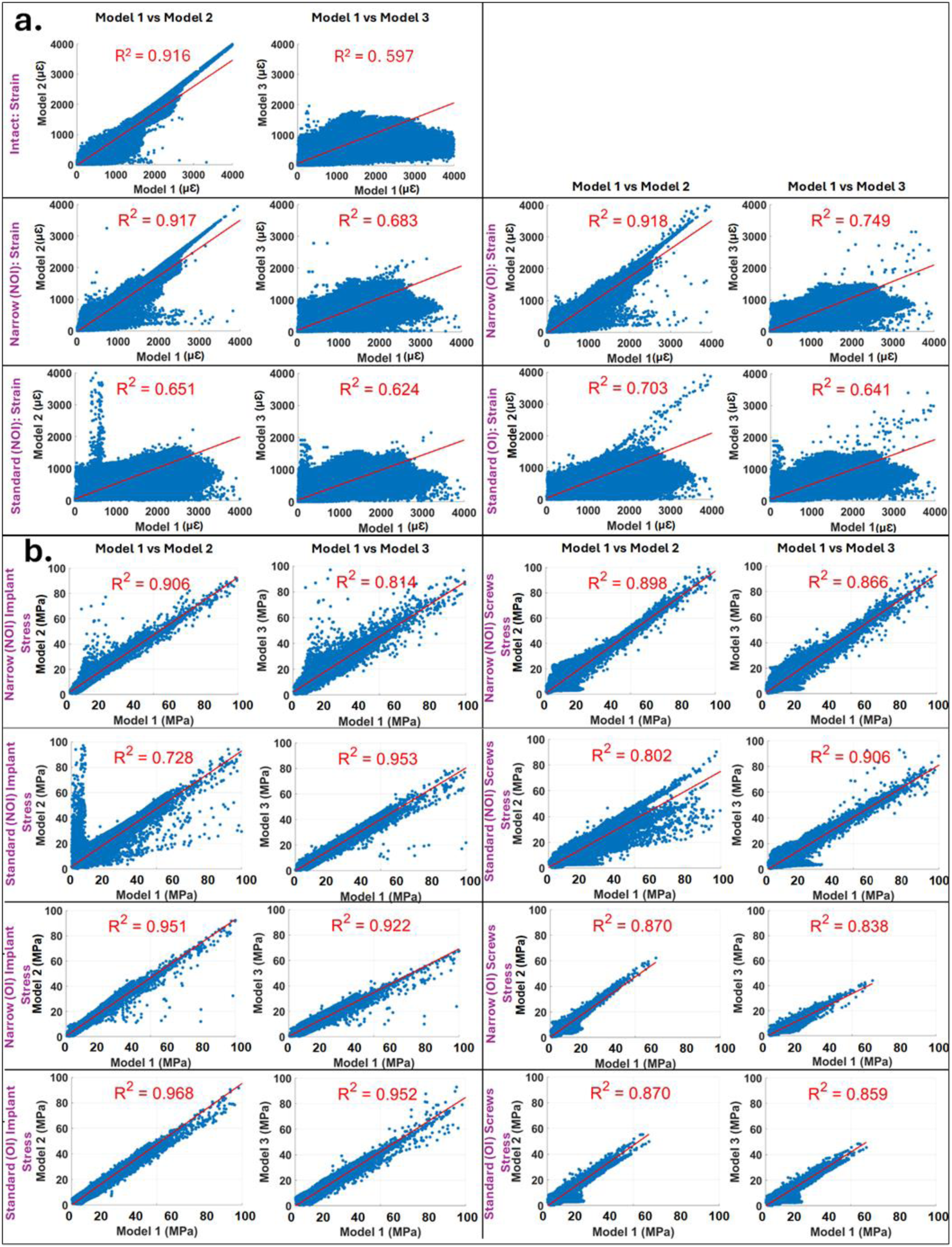
Linear regression plots with coefficient of determination (R2) of FE results for the ICP biting case between the detailed mandible models (Model 1) and the simplified mandible models (Model 2 & 3) in (a) mandible cortical and (b) TMJ implants (mandibular component) and screws.

For the mandibular cortical, R² values ranged from 0.597 to 0.918 for maximum principal strain. Lower R^2^ values were, however, observed primarily owing to the outliers located in the coronoid process, oblique line (near the posterior molar tooth), and interfaces such as the TMJ implant-cortical bone and cortical bone-teeth (especially molars). In addition, the slope of regression line in mandible cortical were in the range 0.486 (with Standard (NOI)) to 0.884 (with Narrow (OI)) between Model 1 & 2, and 0.466 (with Standard (NOI)) to 0.518 (with Narrow (OI)) between Model 1 & 3.

For von-Mises stress in mandibular TMJ implant, R^2^ values ranged from 0.728 to 0.968. In addition, the slope of regression line in mandibular components were in the range 0.910 (Standard (NOI)) to 0.954 (Standard (OI)) between Model 1 & 2, and 0.689 (Narrow (NOI)) to 0.858 (Narrow (NOI)) between Model 1 &3. Similarly, for von-Mises stress in screws, R^2^ values ranged from 0.802 to 0.906. The slope of the regression line in screws was in the range 0.747 (standard (NOI)) to 0.964 (Narrow (NOI)) between Model 1 & 2, and 0.666 (Narrow (OI)) to 0.927 (Narrow (NOI)) between Model 1 &3.

These results indicate that the modelling simplifications, despite capturing the overall strain distribution in the mandibular cortex satisfactorily, influenced the element-specific strain values significantly. However, for TMJ implants and screws, the high R² values and slopes close to unity demonstrate strong agreement between detailed and simplified models for stresses in implants and screws. However, any modelling simplification always resulted in underestimation of stresses in implants and screws as compared to the detailed model.

## Discussion

The present study aims to evaluate the effect of different FE modelling simplifications on the biomechanical assessment of two commercially used mandibular TMJ implants under different clenching tasks. Additionally, the influences of these modelling simplifications on stresses and strains in the mandible, implant, and screws during both non-osseointegrated and osseointegrated conditions have been studied.

An experimental study (Meyer et al., 2002) conducted on an isolated mandible under RMOL bite showed the presence of tensile strains along the anterior border of the ramus and the condylar neck, like the observation made in the presently developed FE models (Fig. 3). The location of the highest principal strain around the oblique line in Model 2 and Model 3 (under INC, ICP, and molar bites) was also in corroboration with the earlier studies(Ghosh et al., 2025; Gröning et al., 2013). Compared to other biting conditions, the cortical bone during the ICP loading showed an extensive region of higher principal strain in the coronoid process and near the intercuspal region across the three modelling techniques. The finding that ICP produced the maximum principal strain on cortical bone among all six clenching tasks was also consistent with the prior investigations (Hijazi et al., 2016; Huang et al., 2015). Additionally, similar to earlier study on dental implants (Niroomand and Arabbeiki, 2020), osseointegration resulted in a reduction in stresses in the implant and screws (Fig. 4).

The current study demonstrates how FE modelling simplifications affect strain distributions in the mandible under various clenching tasks, both during immediate post-operative non-osseointegrated and long-term osseointegrated conditions. A significant difference in the magnitude of strains (up to 50%) between the detailed and simplified mandibles (Fig. 2 and Fig. 3) was observed. Model 2 modelled the entire cancellous bone, teeth, and PDL as that of cortical bone. This assumption caused an increase in the stiffness of the mandible in Model 2 as compared to that of Model 1. Again, Model 3 mandibles were developed from Model 2 mandibles by neglecting the tooth crowns, and, thereby, the mandible stiffness in Model 3 reduced as compared to that of Model 2. In this way, only the detailed mandible models predicted the highest strains under physiological clenching tasks (Fig. 2) and were found to be useful for capturing the risk of mandibular cortex failure.

Similar to the mandible, simplified implanted mandibles were found to reduce von Mises stress in mandibular components by up to 44% overall. Additionally, the simplified mandibles also showed a significant reduction (up to 48%) in von Mises stresses in screws. This clearly indicates an increase in the overall stiffness of the implanted simplified mandible, as explained, resulting in a decrease in stress in the mandibular component and screws. These findings can be corroborated with earlier studies (Beaupied et al., 2007; Currey, 2003; Ghosh et al., 2023; Hart et al., 2017; Jimenez-Palomar et al., 2015) which reveal that a change in the stiffness of the bone modifies the stress-strain behaviour in and around the bone. It should, however, be noted that the overall distribution of stresses and strains in mandibular components and screws was similar between simplified models and the detailed one (Fig. 6; R^2^ up to 0.979).

The present study has a few limitations. The study considered an intact patient-specific mandible model. This study did not consider the effect of dynamic loading conditions while considering multiple modelling simplifications. Although future studies involving dynamic biophysical models with a larger subject cohort might be feasible, the present study provides a thorough insight into the influence of modelling simplifications on preclinical evaluation of TMJ implants. Limitations notwithstanding, the simplified models preserved the overall spatial distribution and qualitative trends of stress and strain in the implants and screws as compared to the detailed mandible FE model. This consistency suggests that, despite underestimating absolute magnitudes, a simplified mandible modelling approach would be useful for preliminary parametric design evaluation of implants and screws. However, a detailed mandible modelling remains essential for finalising the design of TMJ implants and screws.

## Conclusion

The use of simplified models (Model 2 and Model 3) for biomechanical evaluation in finite element studies underestimated the magnitude of maximum principal strain (by ∼16-50%) in cortical bone, while preserving a similar strain distribution. This significant change is attributed to the principle of modelling the entire cancellous bone, teeth, and periodontal ligaments as cortical bone in simplified models. This simplification resulted in stiffening of the overall mandibular FE model. Stresses in implants and screws also exhibited a similar trend, with the detailed model reporting higher stress compared to simplified ones. However, it should be noted that these modelling simplifications did not alter the overall stress-strain distribution in the mandible, implant, and screws during simulated clenching tasks. Thus, these findings suggest that, while simplified mandible models may serve as computationally efficient alternatives for preliminary TMJ implant design, a detailed FE modelling of mandible along with implants and screws should always be considered for final evaluation of TMJ implant designs prior to fabrication and clinical trials.

## CRediT authorship contribution statement

**Girish Chandra:** Conceptualisation, Data Curation, Formal Analysis, Investigation, Methodology, Validation, Visualisation, Writing – original draft. **Rajdeep Ghosh:** Conceptualisation, Formal Analysis, Investigation, Methodology, Software, Visualisation, Writing – original draft. Writing – review & editing. **Vivek Verma:** Data Curation, Formal Analysis, Investigation, Visualisation. **Kamalpreet Kaur:** Data Curation, Formal Analysis, Investigation, Visualisation. **Ajoy Roychoudhury:** Funding acquisition, Project Administration, Resources, Supervision. **Sudipto Mukherjee:** Conceptualisation, Funding acquisition, Methodology, Project Administration, Resources, Supervision, Writing – review & editing. **Anoop Chawla:** Conceptualisation, Funding acquisition, Methodology, Project Administration, Resources, Supervision, Writing – review & editing. **Kaushik Mukherjee:** Conceptualisation, Funding acquisition, Methodology, Project Administration, Resources, Supervision, Writing – review & editing.

## Acknowledgments

The study was supported by the Indo–German Science & Technology Centre (IGSTC), under grant no. IGSTC/Call 2020/add-bite/48/2021-22/260.

## Figure Legends

**Supplementary Figure 1.**
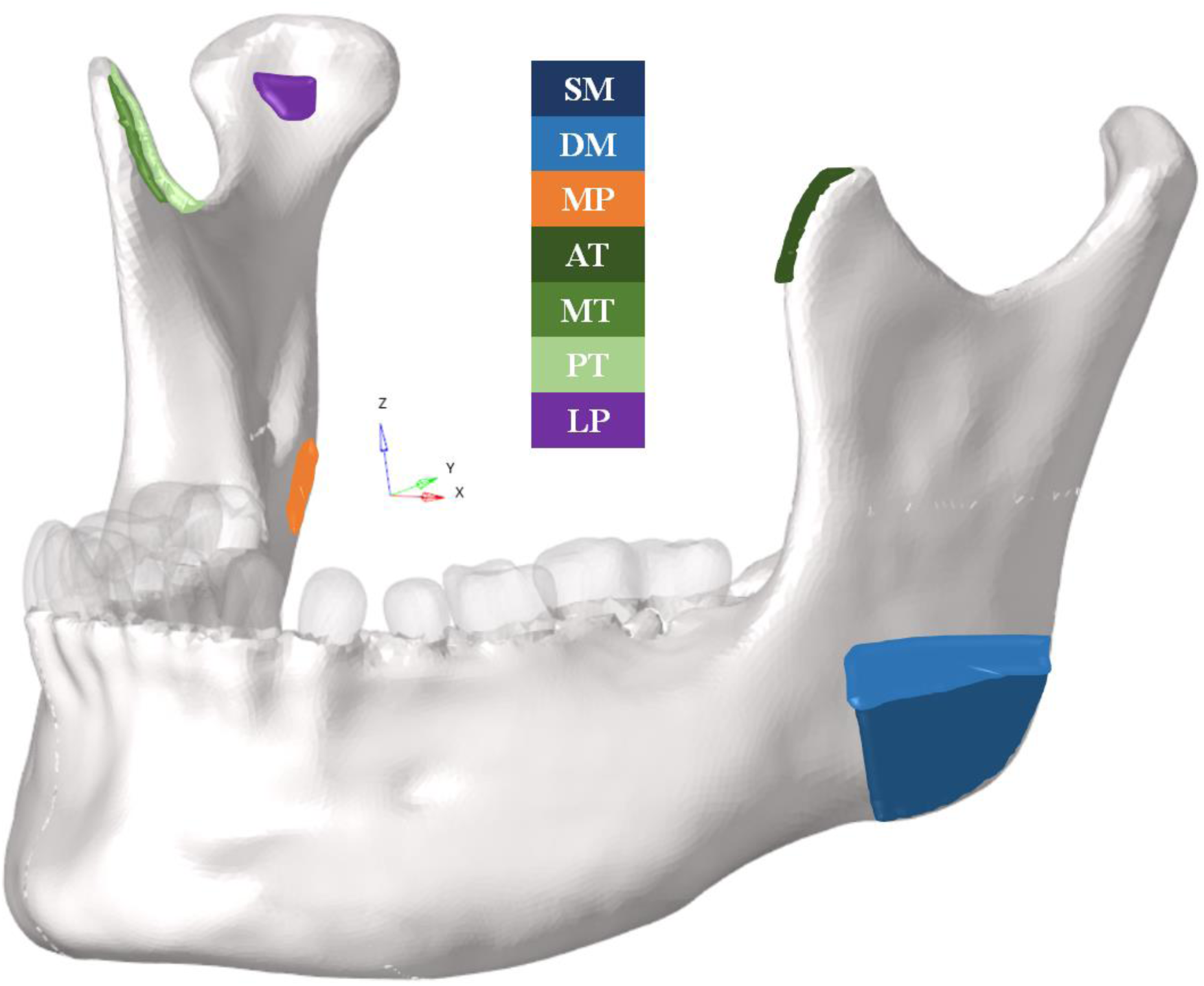
Different muscle locations (SM: Superior Masseter, DM: Deep Masseter, MP: Medial Pterygoid, AT: Anterior Temporalis, MT: Middle Temporalis, PT: Posterior Temporalis, LP: Lateral Pterygoid) considered in the FE models

**Supplementary Figure 2:**
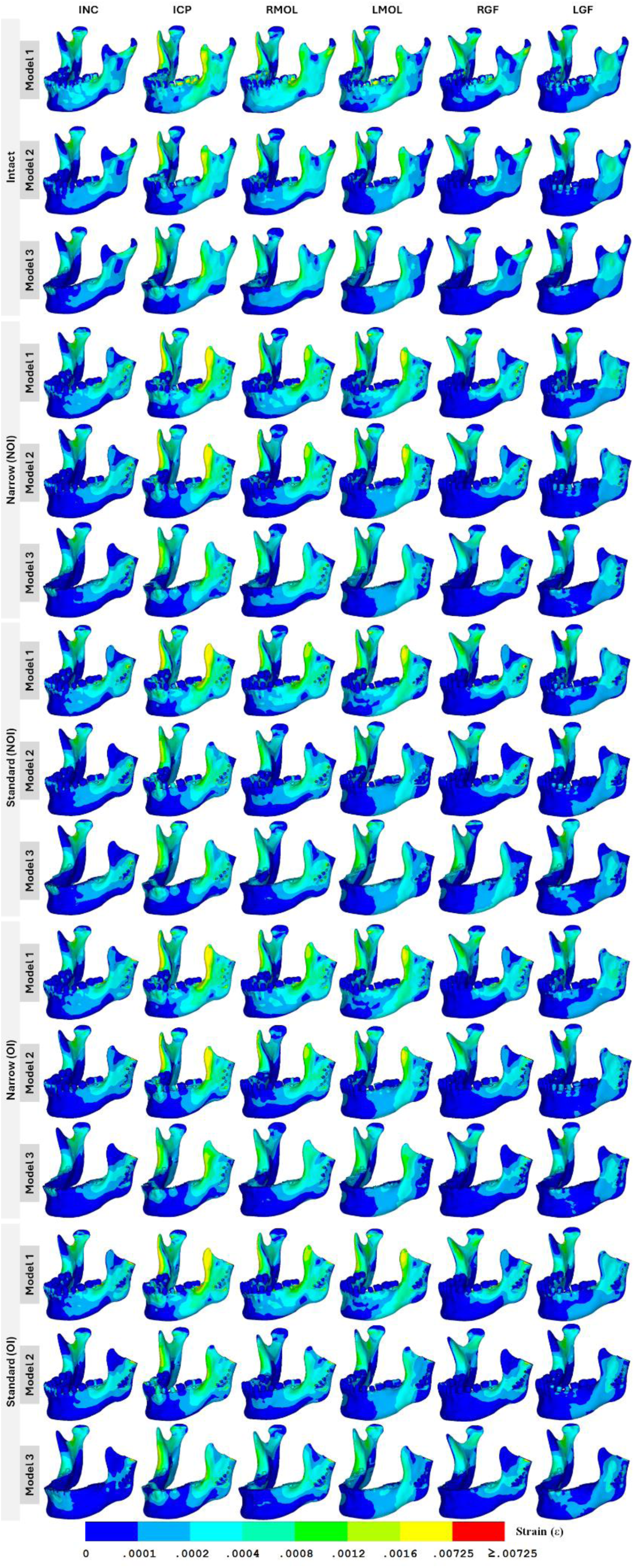
Comparison of maximum principal strain distribution on the cortical bone in various FE models (Model 1: Detailed Segmented Mandible, Model 2: Cortical Mandible, Model 3: Cortical mandible without crown of teeth, NOI: non-osseointegrated, OI: osseointegrated)

**Supplementary Figure 3:**
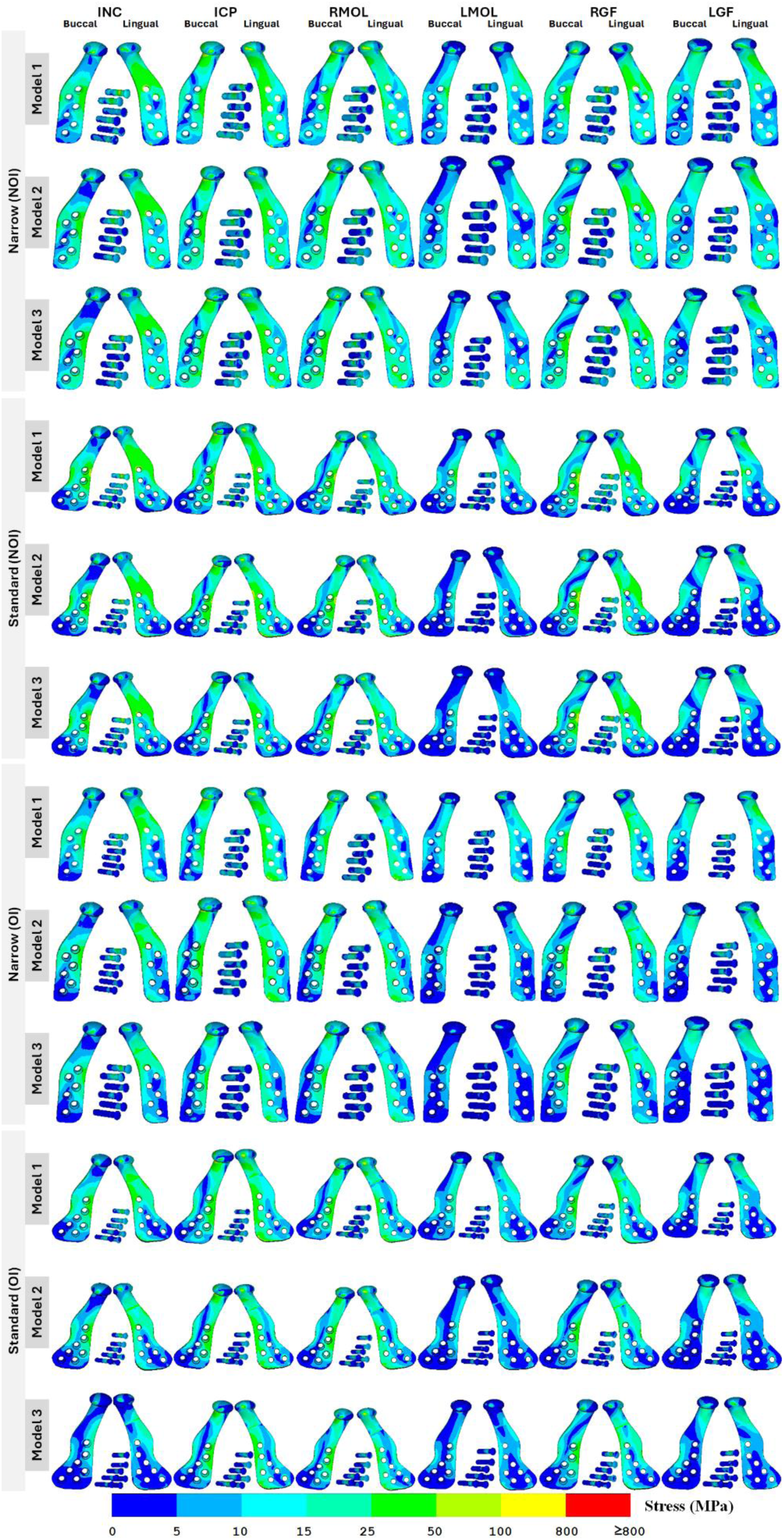
Comparison of maximum von Mises stress distribution on the TMJ implants (Mandibular component and Screws) in various FE models (Model 1: Detailed Segmented Mandible, Model 2: Cortical Mandible, Model 3: Cortical mandible without crown of teeth, NOI: non-osseointegrated, OI: osseointegrated)

## List of Tables

**Supplementary Table 1.**
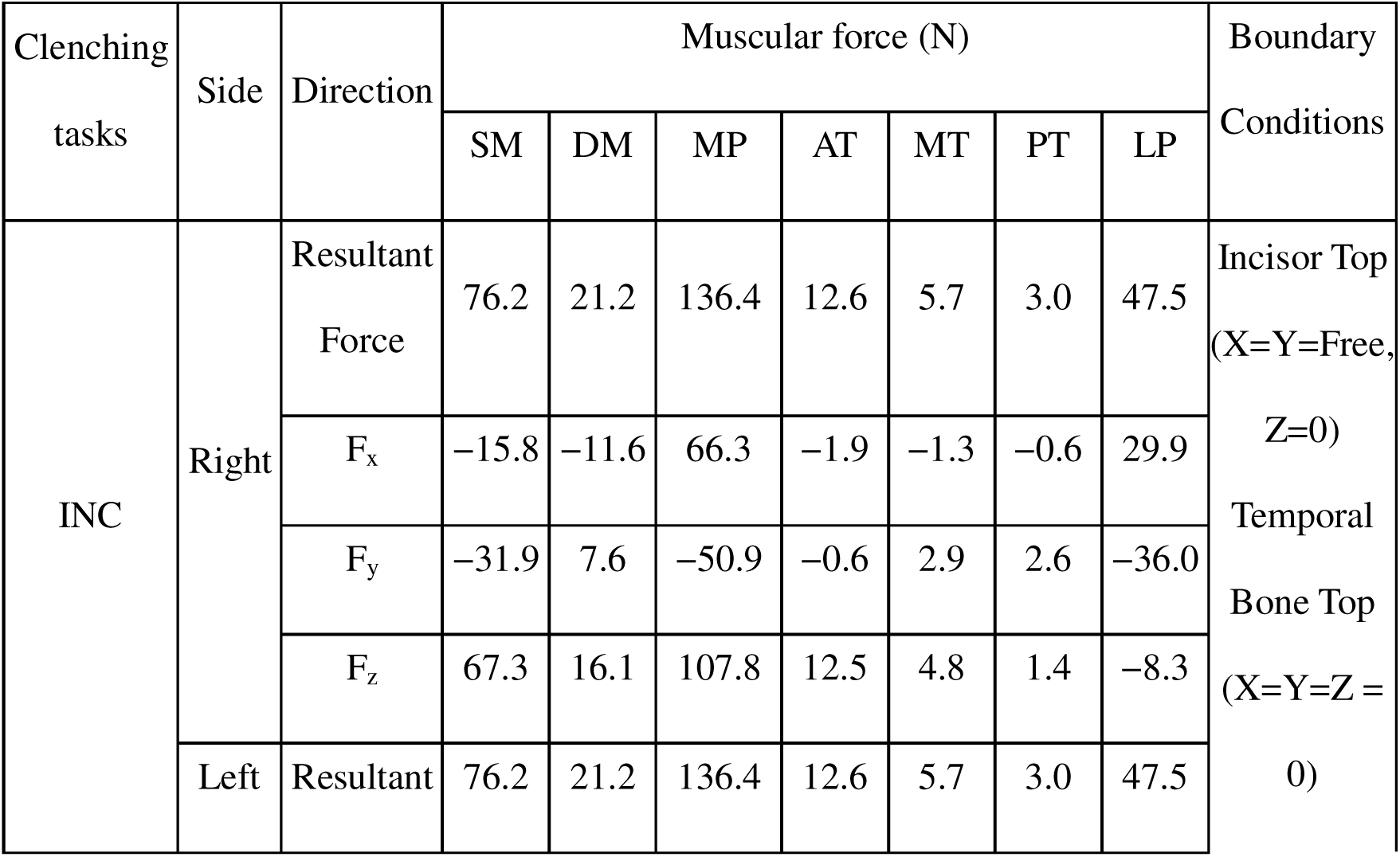
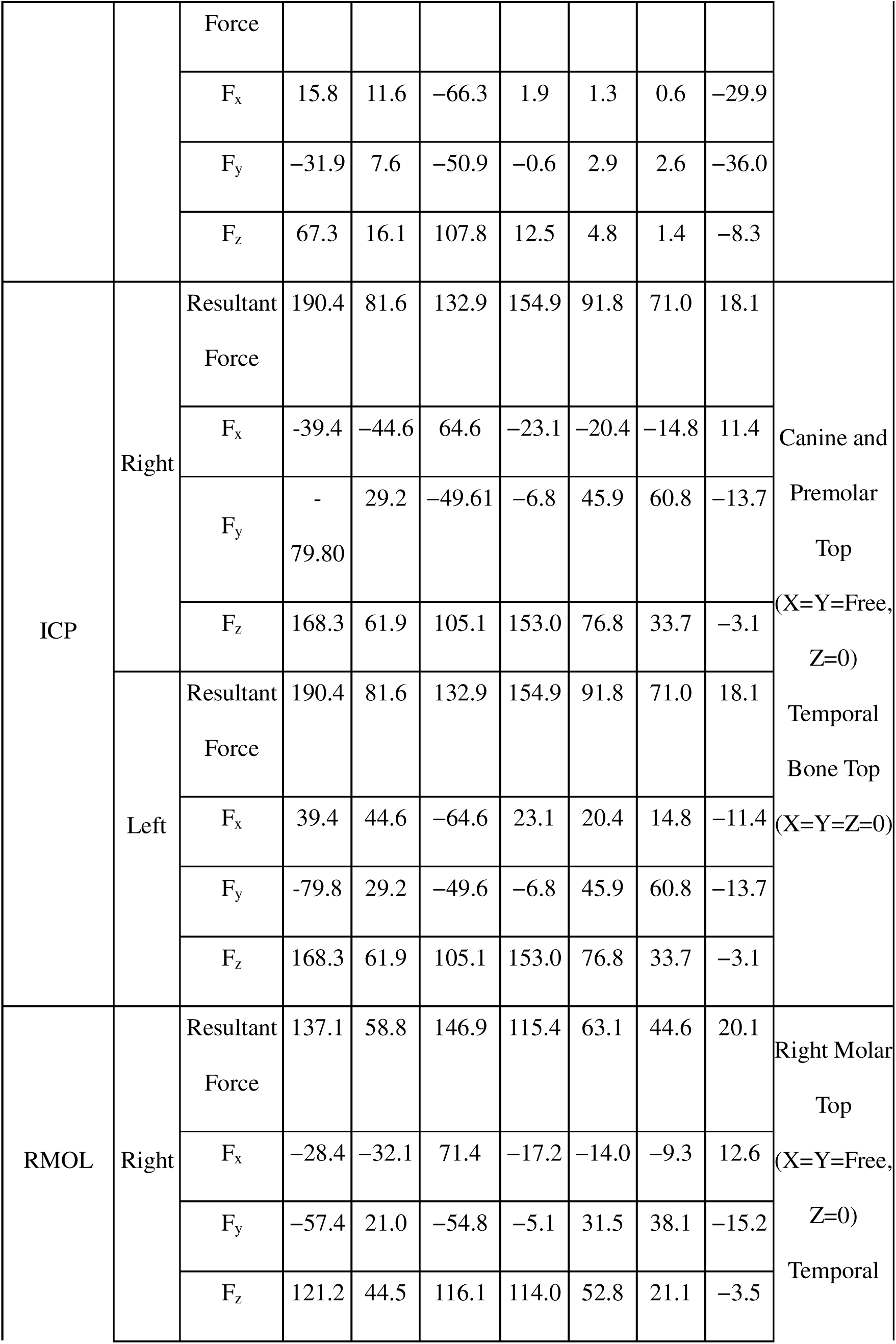
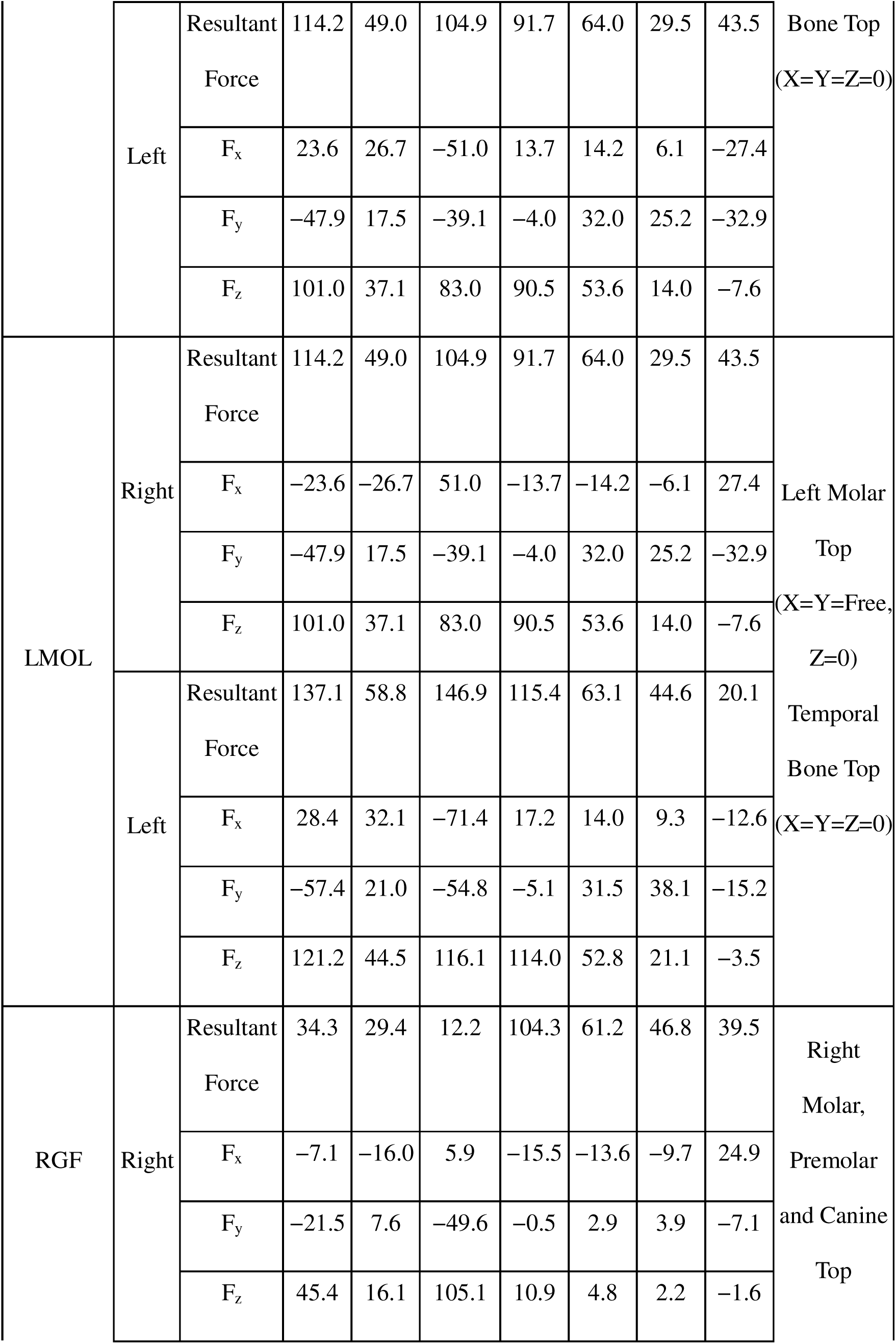
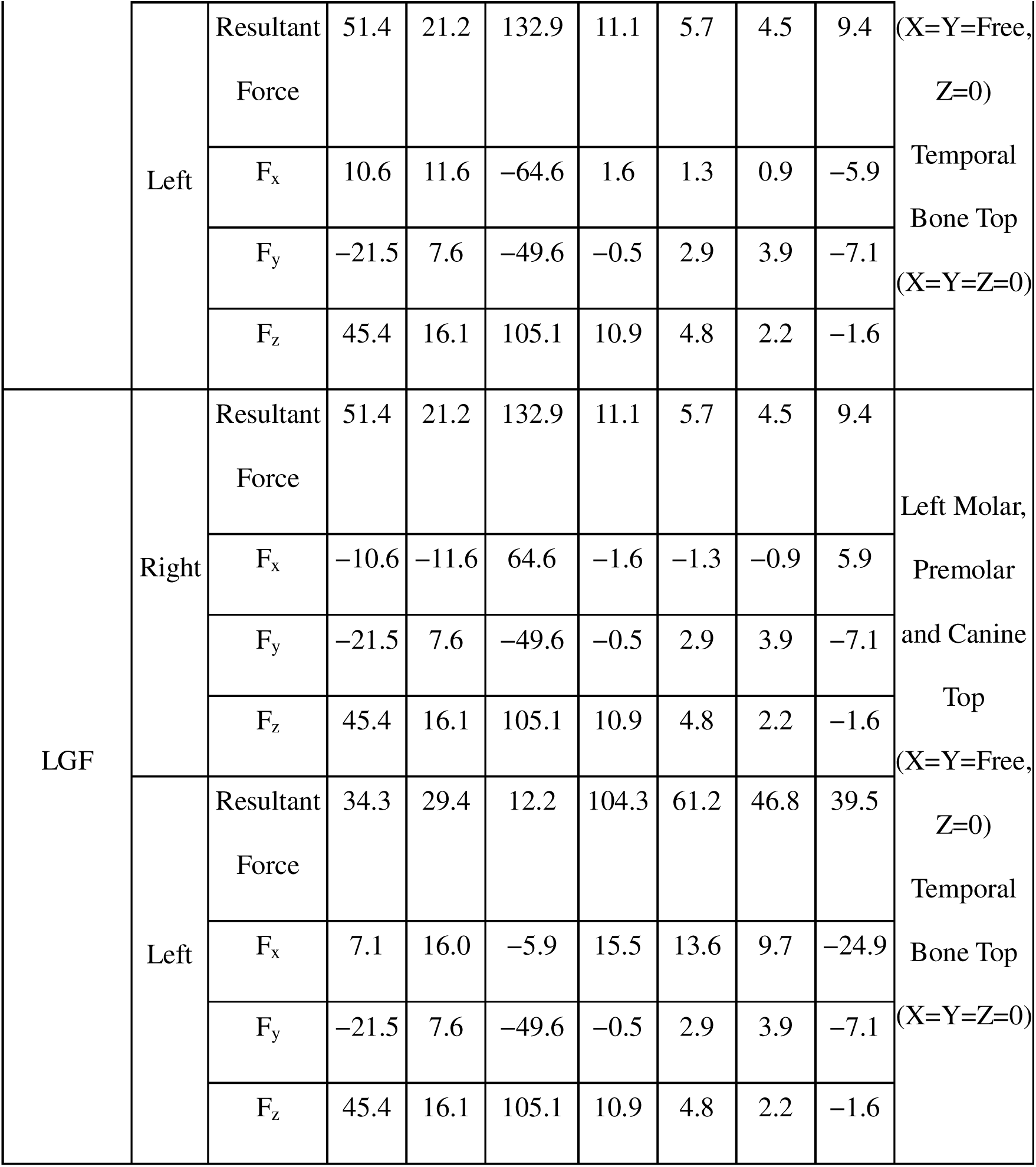
Muscular force and boundary conditions of clenching activities (Dutta et al., 2023; Hijazi et al., 2016).

